# Singular Value Decomposition for the Automated Analysis of NMR Fraction Libraries

**DOI:** 10.64898/2026.10.30.763183

**Authors:** Christopher Esselman, Arthur S Edison, James H Prestegard

## Abstract

Identifying and quantifying molecular components of biological fluids offers great potential for the diagnosis of disease, detecting environmental contaminants, and process monitoring. The use of simple 1D ^1^H NMR spectra to accomplish this task offers advantages in terms of broad applicability and quantitative output. However, it also encounters obstacles in extracting individual components from complex superimposed spectra and in compound identification through comparison with limited database entries. Here, we present a new protocol that includes automatically separating component spectra using singular value decomposition (SVD) and matching them to database entries that carry both chemical shift and scalar coupling information. The procedure is illustrated by application to a fungal extract from *Neurospora crassa* that is chromatographically fractionated to form a two-dimensional spectral fraction library. Results are compared to a previous analysis that used a semi-manual curation of separated spectra followed by chemical shift matching.

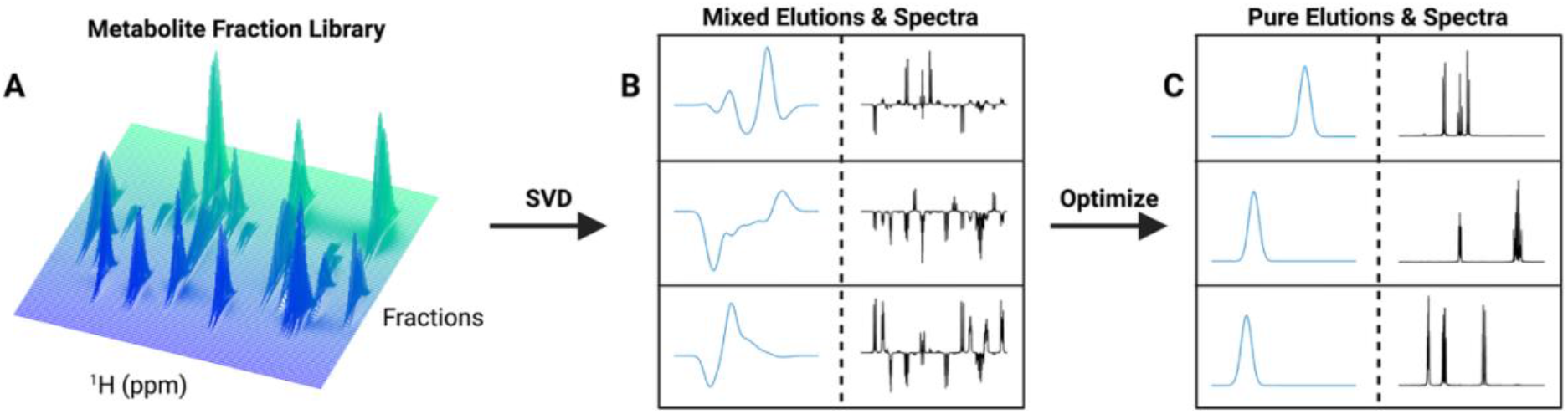

## INTRODUCTION

One-dimensional proton NMR spectroscopy (1D ^1^H NMR) offers great potential for monitoring and control of biological processes (*1*). The spectra are highly quantitative, and the ubiquity of protons (hydrogens) in metabolic products provides dispersed resonances from which the identity and concentration of metabolites can be derived. Perhaps the best example of impact on the biomedical field is the now common use of 1D ^1^H NMR to provide apolipoprotein profiles of blood samples. Between 2018 and 2024, a single provider logged more than 1 million analyses (*2*). The connection between these profiles and cardiac health is well established (*3*), but further analysis of the underlying spectra has suggested value in diagnosing diseases ranging from diabetes to long COVID (*4, 5*). There is also growing interest in using 1D ^1^H NMR to optimize disease treatment once a diagnosis is made (*6*), and beyond the medical field, there are opportunities to use metabolic signatures to optimize cell cultures for the production of materials and the detection of environmental hazards (*7, 8*). An extensive review of these and other applications of NMR-based metabolomics has recently appeared (*9*). Even with these numerous examples, the potential is far from fully realized. The overlap of peaks from hundreds of metabolites normally present in biological fluids often obscures valuable correlations and prevents the extraction of a sufficient spectral segment for metabolite identification.

There have been efforts to improve resolution and expand compound identification by adding additional dimensions to spectral acquisition (*10*). However, there is usually some loss of quantitation ability, and resolution is always limited. Hence, exploration of additional approaches to extracting individual metabolite spectra is worthwhile. We have taken some steps toward this exploration by generating a chromatographic “metabolite fraction library” (mFL) in which we separated an extract from *Neurospora crassa* into more than 100 fractions and collected individual spectra on each (*11*). Using a tool we call “FLAnalyzer”, spectral peaks were then grouped via correlation to assemble a “metabolite basis set” (mBS), and compound identification was achieved by matching chemical shifts of each basis element to a chemical shift database.

Currently, generating an mBS is time-consuming and requires substantial manual intervention. Furthermore, database matching that uses only chemical shifts fails to fully exploit the multiplet structure and multiplet connections present in a complete spectrum.

Here, we explore the use of singular value decomposition (SVD) to automate the extraction of individual metabolite spectra from our mFL. SVD is a well-documented approach to reducing the dimensions of data matrices (*12, 13*). More specifically, in this application, we reduce our mFL matrix to three matrices: an elution vector matrix with **k** elements, a **k** x **k** singular matrix, and a spectral vector matrix with **k** elements. The singular matrix is diagonal, with elements ordered by importance. The size of these elements drops abruptly at certain values of **k**, providing an estimate of the number of metabolites having significant concentrations.

Furthermore, truncation at selected values of **k** drastically reduces the dataset size, effectively filtering out baseline and spectral noise.

While the determination of the number of major metabolites and reduction in dimensions are useful, the vectors in the initial elution and spectral matrices do not initially represent properties of a single compound. However, the vectors do form orthonormal sets. This allows us to use standard optimization routines to find linear combinations representing single compounds. For elution vectors, optimization was based on minimizing the difference between the linear combination of SVD components and an experimentally derived target function. Optimal coefficients for SVD components were collected into a transformation matrix, which was then applied to both the initial elution and spectral sets. The spectral vectors were then further optimized using a target function that minimized error between a linear combination of spectral vectors and a covariance spectrum derived from the primary elements of each elution vector.

Optimal coefficients were then assembled into a new elution transformation matrix, and the transformation process was repeated until convergence to a single transformation matrix was achieved. The procedure, implemented in MATLAB, is outlined in Figure 1 and described more fully in the methods section of this paper.

**Figure 1.**
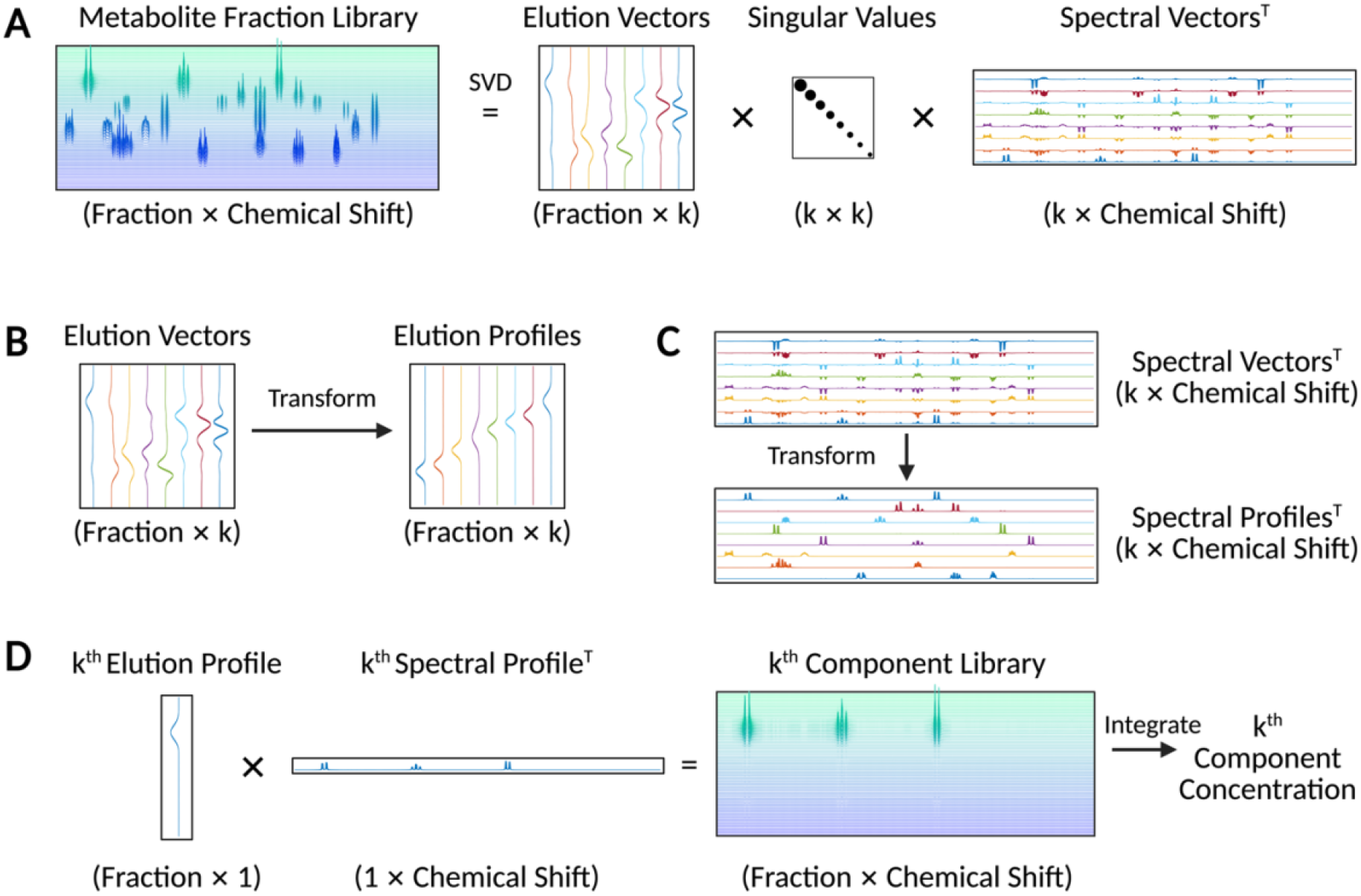
Flow chart for decomposition of a fraction library by SVD. (**A**) Initial SVD decomposition. (**B**) Transformation of elution vectors to elution profiles of individual compounds. (**C**) Transformation of spectral vectors to spectra of individual compounds. (**D**) Outer products of elution and spectral profiles generate single compound maps, and their integration yields concentrations.

Once spectral vectors are obtained, compound identification can be pursued. As the SVD procedure returns not a list of peaks, but an entire spectrum containing multiplet structures and distortions from strong-coupling effects, it makes sense to match the entire transformed spectrum to a spectrum in a metabolite database. Fortunately, entire spectra collected (or simulated) across all commonly used magnetic fields are available in databases such as GISSMO (*14*) and the HMDB (*15*). We have combined spectra from these databases and, using chemical shifts and coupling constants from the literature, added spectra, particularly some from carbohydrates (*16, 17*). As our extracted spectra have specific line shapes spanning a fixed number of points, all spectra in our assembled database were transformed to have similar spectral properties and normalized. Searching for matches can then be done using a simple dot-product score (equivalent to cosine similarity) that ranges from 1.0 (perfect match) to 0 (no match). This procedure has been implemented in MATLAB and is described more fully in the methods section.

An additional advantage of the SVD approach is that we can reconstruct the original library from the product of the elution profiles, singular values, and spectral components. Any single product gives a matrix that can be summed (integrated) to give a relative concentration. By reconstructing the initial library, we can also assess how well that library is reproduced by our purified spectral and elution profiles, providing a quantitative measure of completeness of the fraction library analysis. Using a single iteration (**k** = 58) of our SVD approach, we captured 98.26% of the variance in the *N. crassa* library (Figure S1). Removal of contributors to a reconstruction at any **k** value also allows us to better observe residual metabolites not included in the reconstruction.

## RESULTS

### Deconvolution of Peaks in the Fraction Library

To illustrate the ability of the SVD approach to handle severe overlap of elution peaks, we compare our SVD method with one based solely on correlation analysis (Figure 2) (*11*). This example uses a spectral vector eventually assigned to histidine. While grouping peaks by correlation across eluted fractions works well for well-resolved signals, it struggles when distinct compounds co-elute and overlap. In the covariance spectrum, selecting the highly correlated components (red) successfully retrieves the histidine multiplets at 7.85, 7.10, 3.95, and 3.10 ppm, but fails to capture the multiplet at 3.25 ppm, which is obscured by an overlapping signal. In contrast, the SVD reconstruction leverages orthogonal differences in chromatographic elution profiles across the two-dimensional data matrix to mathematically separate the co-eluting signals. As a result, the SVD method accurately models the overlapped multiplet, yielding a purified spectral vector suitable for automated database matching.

**Figure 2.**
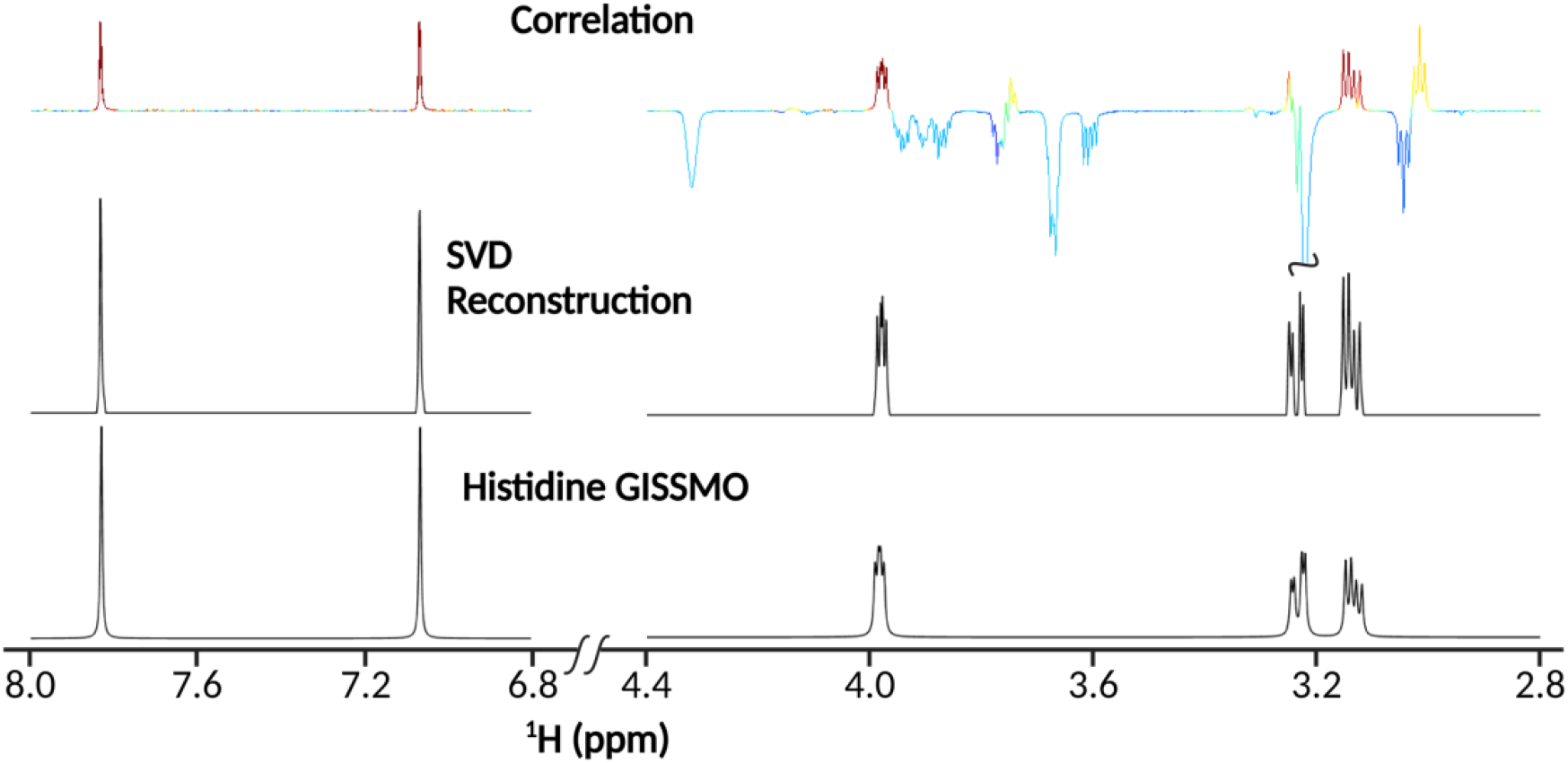
SVD optimization deconvolutes overlapping peaks. (**Top Spectrum**) The covariance spectrum using the histidine peak at ∼ 7.1 as a driver in the *N. crassa* mFL. The spectrum was calculated using just the fractions in which histidine eluted. Color indicates the correlation between the peaks and the driver, with highly correlated peaks shown in red. (**Middle Spectrum**) The spectral vector of histidine created automatically using the SVD optimization algorithm. (**Bottom Spectrum**) The histidine GISSMO spectrum.

### Database Matching Spectral Vectors Directly

To further illustrate our method, we focus on the segment of our fraction library depicted in Figure 3A. This segment (fractions 79 to 131 of 140) contains some of the most intense peaks in the library, as well as peaks falling within a highly congested region (3.0 to 4.0 ppm). Congestion presents a problem for more traditional analysis methods, which rely on peak picking followed by queries against text-based databases such as COLMAR or the BMRB (*18, 19*). Highly crowded spectral regions dominated by multiplets rather than single peaks produce substantial overlap, leading to ambiguous or false-positive assignments. Our proposed approach reduces ambiguity by shifting the matching from discrete chemical-shift coordinates to continuous spectral profiles. This preserves crucial structural information, including relative peak intensities and J-coupling information. This is illustrated in Figure 3B, which focuses on four spectral vectors isolated using our SVD approach. Two of the elements have more than 20 peaks in the 3.0 to 4.0 spectral region. Figure 3B shows the cosine similarity metric for each element. A clear distinction among the four elements is made, assigning element 2 to trehalose. Trehalose is a symmetric α-linked dimer of glucose with only 7 distinct chemical shifts. While the presence or absence of peaks outside the 3.0 to 4.0 ppm region would help in assignment, it is clear that assignments based purely on chemical shifts would have to discount the absence of numerous matches in the 3.0 to 4.0 ppm region in order to make an unambiguous assignment.

**Figure 3.**
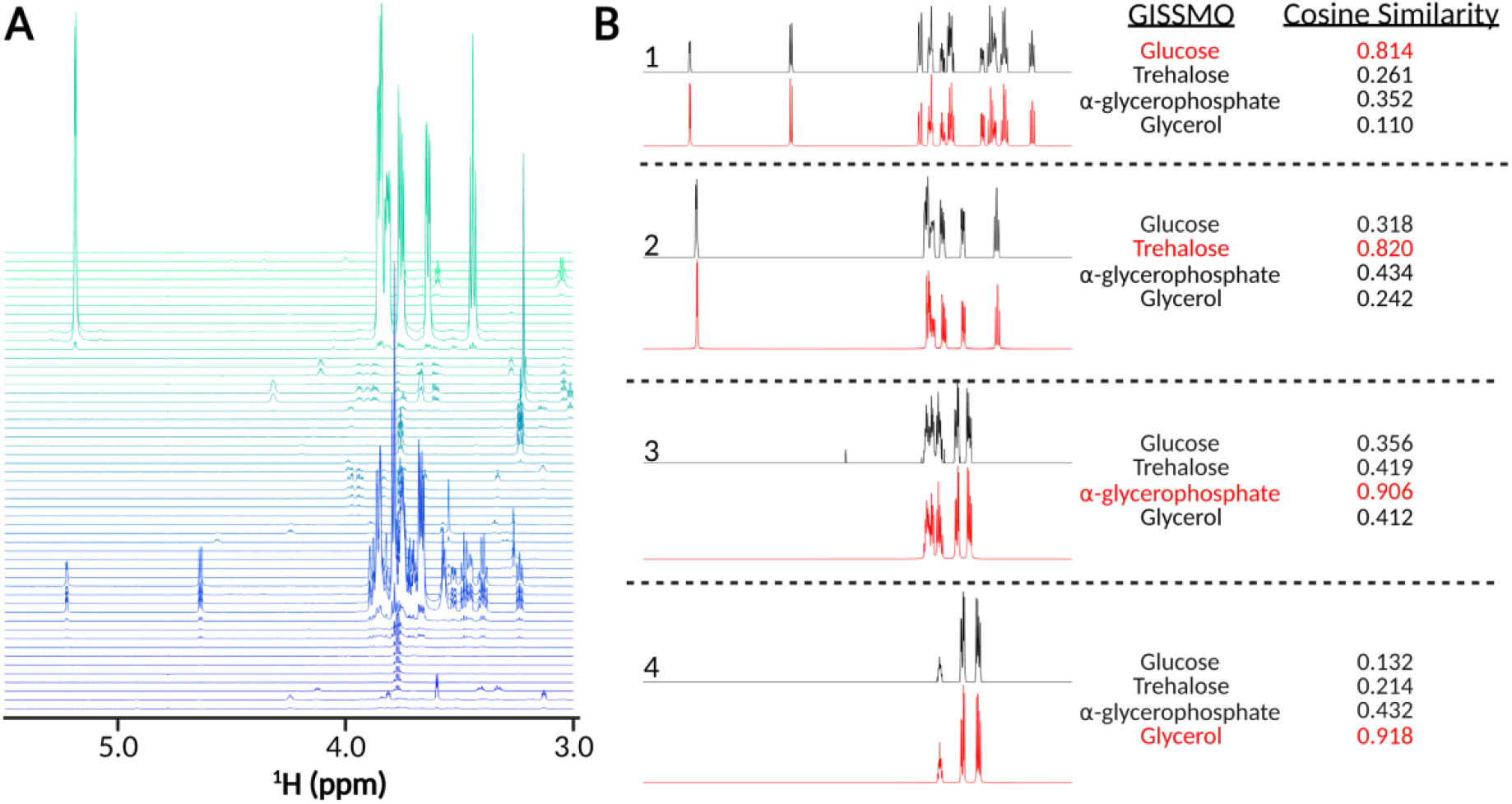
The SVD approach distinguishes compounds with similar chemical shifts. **(A)**. A region of the *N. crassa* mFL containing some of the most intense and overlapped peaks. **(B)** Cosine similarity scoring of SVD basis set elements against the GISSMO database. The four SVD basis set elements are shown in black, along with the best-matching GISSMO reference spectra shown in red. In each case, four distinct cosine similarity scores are presented.

Table 1 summarizes the compound assignments and cosine similarity scores for the spectral vectors extracted from the *N. crassa* mFL segment in Figure 2A. Direct spectral comparison between the fully automated SVD reconstructions and the semi-automated FLAnalyzer-derived basis set confirms that SVD reliably recovers the underlying metabolite signatures across this congested region, yielding a median similarity score of 0.948, with 19 of the 33 vectors exceeding 0.90. 17 SVD vectors achieved GISSMO similarity scores above 0.80 (0.811–0.954), and 16 of the top 19 elements (GISSMO scores > 0.696) produced automated GISSMO assignments identical to the FLAnalyzer annotations.

**Table 1.**
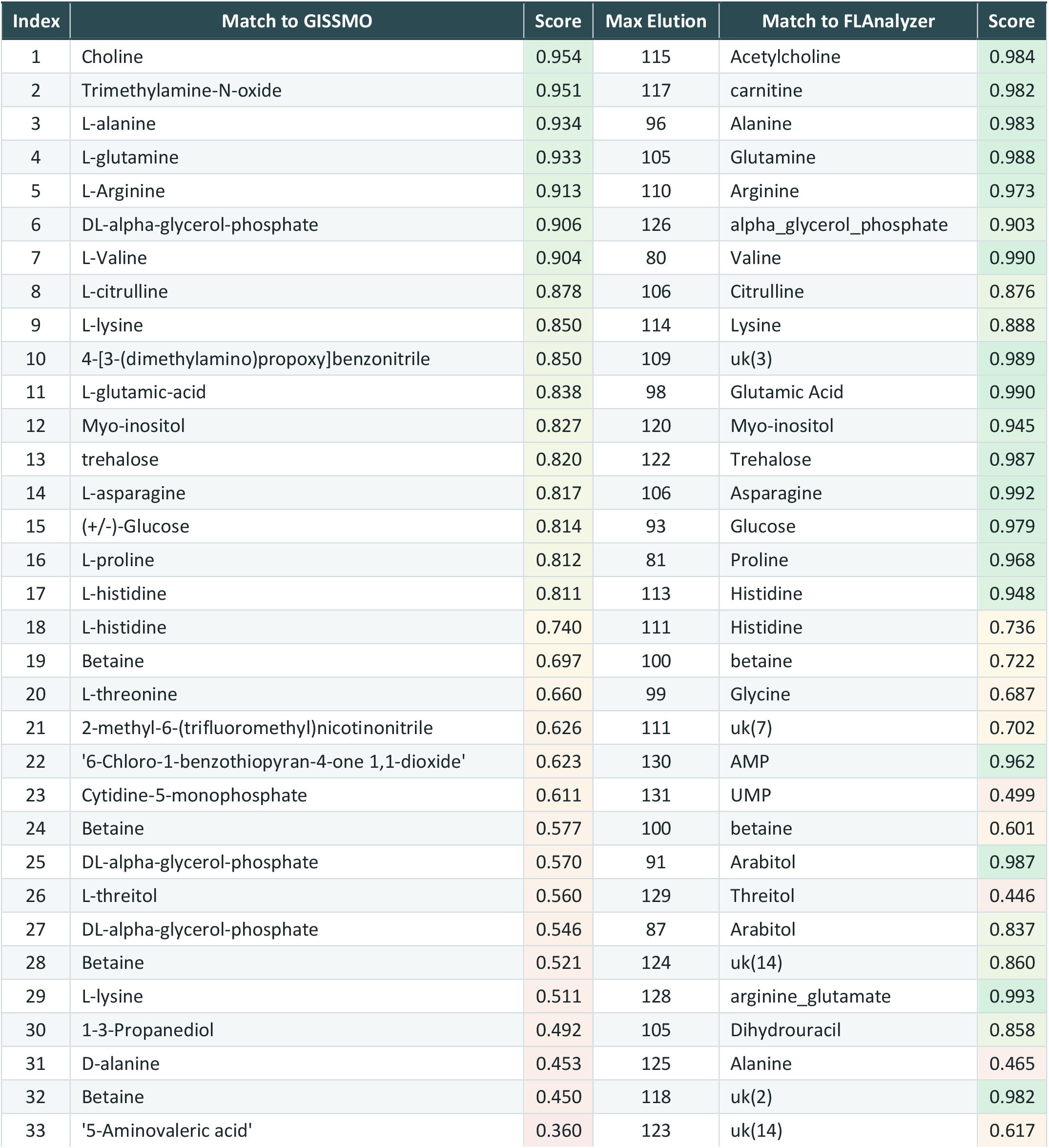
Database Matching Spectral Vectors.

| Index | Match to GISSMO | Score | Max Elution | Match to FLAnalyzer | Score |
|---|---|---|---|---|---|
| 1 | Choline | 0.954 | 115 | Acetylcholine | 0.984 |
| 2 | Trimethylamine-N-oxide | 0.951 | 117 | carnitine | 0.982 |
| 3 | L-alanine | 0.934 | 96 | Alanine | 0.983 |
| 4 | L-glutamine | 0.933 | 105 | Glutamine | 0.988 |
| 5 | L-Arginine | 0.913 | 110 | Arginine | 0.973 |
| 6 | DL-alpha-glycerol-phosphate | 0.906 | 126 | alpha_glycerol_phosphate | 0.903 |
| 7 | L-Valine | 0.904 | 80 | Valine | 0.990 |
| 8 | L-citrulline | 0.878 | 106 | Citrulline | 0.876 |
| 9 | L-lysine | 0.850 | 114 | Lysine | 0.888 |
| 10 | 4-[3-(dimethylamino)propoxy]benzonitrile | 0.850 | 109 | uk(3) | 0.989 |
| 11 | L-glutamic-acid | 0.838 | 98 | Glutamic Acid | 0.990 |
| 12 | Myo-inositol | 0.827 | 120 | Myo-inositol | 0.945 |
| 13 | trehalose | 0.820 | 122 | Trehalose | 0.987 |
| 14 | L-asparagine | 0.817 | 106 | Asparagine | 0.992 |
| 15 | (+/-)-Glucose | 0.814 | 93 | Glucose | 0.979 |
| 16 | L-proline | 0.812 | 81 | Proline | 0.968 |
| 17 | L-histidine | 0.811 | 113 | Histidine | 0.948 |
| 18 | L-histidine | 0.740 | 111 | Histidine | 0.736 |
| 19 | Betaine | 0.697 | 100 | betaine | 0.722 |
| 20 | L-threonine | 0.660 | 99 | Glycine | 0.687 |
| 21 | 2-methyl-6-(trifluoromethyl)nicotinonitrile | 0.626 | 111 | uk(7) | 0.702 |
| 22 | '6-Chloro-1-benzothiopyran-4-one 1,1-dioxide' | 0.623 | 130 | AMP | 0.962 |
| 23 | Cytidine-5-monophosphate | 0.611 | 131 | UMP | 0.499 |
| 24 | Betaine | 0.577 | 100 | betaine | 0.601 |
| 25 | DL-alpha-glycerol-phosphate | 0.570 | 91 | Arabitol | 0.987 |
| 26 | L-threitol | 0.560 | 129 | Threitol | 0.446 |
| 27 | DL-alpha-glycerol-phosphate | 0.546 | 87 | Arabitol | 0.837 |
| 28 | Betaine | 0.521 | 124 | uk(14) | 0.860 |
| 29 | L-lysine | 0.511 | 128 | arginine_glutamate | 0.993 |
| 30 | 1-3-Propanediol | 0.492 | 105 | Dihydrouracil | 0.858 |
| 31 | D-alanine | 0.453 | 125 | Alanine | 0.465 |
| 32 | Betaine | 0.450 | 118 | uk(2) | 0.982 |
| 33 | '5-Aminovaleric acid' | 0.360 | 123 | uk(14) | 0.617 |

Discrepancies between the automated GISSMO hits and the FLAnalyzer annotations fall into two clear categories. First, elements 1 and 2 were matched to acetylcholine (0.984) and carnitine (0.982) using manual annotation, whereas automated GISSMO queries returned structurally related quaternary amines choline (0.954) and trimethylamine-N-oxide (0.951). In both cases, the assignment is complicated by the dominance of the spectra by the trimethylamine peak. It is a singlet with an intensity corresponding to 9 equivalent protons. Manual examination of the spectra raises questions about the acetylcholine assignment, as it lacks intensity in the regions expected for the acetyl methyl group. Also, choline is not likely to be correct, as there is a significant mismatch for the chemical shift of the triplet expected for a methylene next to the hydroxyl group of choline. This is likely a choline derivative absent in current databases. A suggested assignment is presented in the discussion. Second, some SVD elements matched FLAnalyzer with high fidelity (>0.85) but lacked a corresponding high-scoring match in GISSMO. For instance, elements 25 (arabitol) and 32 (uk2) exhibited near-unity agreement with the FLAnalyzer basis spectra (0.987 and 0.982, respectively) despite lower GISSMO scores (0.570 and 0.450), effectively flagging either compound as outside our expanded GISSMO database. In fact, arabitol is not in our database.

## Automatic Integration of Compounds in the Fraction Library

In Figure 4, we evaluate the SVD approach’s ability to quantify compound concentrations directly from the mFL by comparing automated results with those obtained by manual integration. Manual quantification requires an analyst to isolate individual multiplets, integrate them across fractions, sum over the elution profile, and reference the total area to an internal standard (DSS). In contrast, the SVD framework automates this calculation by integrating the outer product of the elution and spectral vectors, multiplying by the associated singular value, and dividing by the number of integrable protons in the assigned compound to determine concentrations relative to the DSS reference signal. As shown in Figure 4, the SVD-derived concentrations exhibit a strong correlation with manual integrations over a wide dynamic range. An R^2^ of 0.923 was calculated as 1 minus the ratio of the sum of squared residuals to the total sum of squares. This confirms the reliability of automated quantification across a wide range of library concentrations.

**Figure 4.**
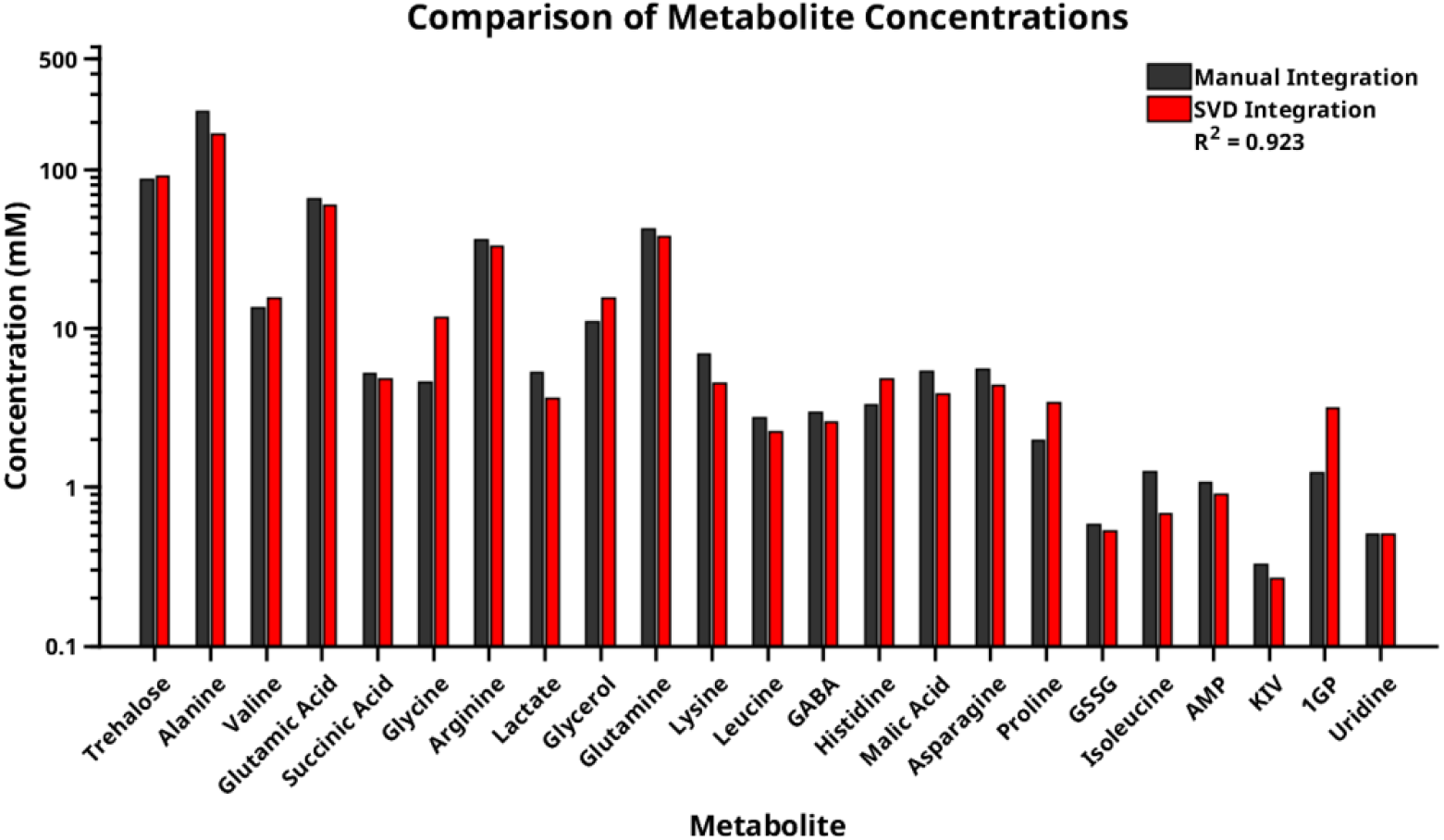
SVD automatically integrates compounds in the mFL. Concentration of metabolites from manual integration (black bars) versus SVD automated integration (red bars) of annotated metabolites within the *N. crassa* mFL.

## DISCUSSION

We believe that the above results demonstrate that Singular Value Decomposition (SVD), when combined with covariance processing of target spectra, provides an effective, automated approach for deconvoluting highly overlapped one-dimensional ^1^H NMR metabolite fraction libraries. By distilling complex matrices into distinct elution and spectral vectors, SVD generates pure, full-multiplet spectra that represent individual metabolites. These isolated spectra can be directly queried against established databases for structural identification. Furthermore, even when an extracted spectrum does not yield a high-scoring database match, it represents a mathematically isolated, physical component. Therefore, these unidentified pure spectra can serve as spectral vectors for the quantitative fitting and monitoring of data in broader metabolomic studies.

Moreover, SVD’s ability to extract the entire compound spectrum, complete with multiplet structures, is highly advantageous for database matching. Many traditional NMR metabolomics workflows rely solely on chemical-shift peak picking, an approach that discards structural information from scalar couplings and relative peak intensities. This loss of information is especially detrimental to carbohydrate identification, which is hampered by severe spectral crowding and complex multiplet overlaps. By matching the full spectrum against repositories such as the GISSMO database, we retain this vital data. To maximize this capability, we supplemented the GISSMO database with additional simulated carbohydrate spectra, allowing us to identify several monomers, dimers, and anomeric mixtures confidently.

Furthermore, paired spectral and elution profiles provide a significant advantage for quantitative analysis by allowing us to integrate metabolites directly from the metabolite fraction library rather than relying on manual integration. Finally, the ability to reconstruct the original library from the product of the elution profiles, singular values, and spectral components of major species systematically isolates a residual matrix (Figure S2). This allows us to observe lower-concentration metabolites by physically isolating them, so they can be parsed out in subsequent steps rather than being lost in the noise.

It is not our intention here to go beyond quantifying the presence of major *N. crassa* metabolites and assessing the reasons for their abundance. However, some of the strongest peaks in the segment highlighted in Figure 2A deserve some comment. The strongest peaks are assigned to trehalose, a symmetric non-reducing carbohydrate, formed by linking the anomeric carbons of two glucose residues in an α configuration. Trehalose is one of the many carbohydrates missing from the initial GISSMO database. We added it by matching the 700 MHz spectrum from the BMRB using Spinach (*17*) and then simulating an 800 MHz spectrum using the extracted chemical shifts and coupling constants. While this process is tedious, there is now a database (CASMDB) that allows inclusion of a “matrix.multiplet” in the NEF format (NMR Exchange Format) (*20*). This matrix contains the chemical shift and coupling constant data needed for spectral simulation at any field strength. Encouraging deposition in this format could streamline the process. But why is trehalose synthesized in such abundance by *N. crassa*? This is a question with a probable answer. Trehalose is a known osmolyte and protectant against the effects of dehydration (*21*).

A good match for the second-strongest peak could not be found in any standard database, but it was clearly related to choline. An examination of the literature on fungal organisms suggested that choline sulfate may be a better match for this peak. Sulfation produces no additional proton peaks in the spectrum but will change the chemical shift of the methylene triplet next to the hydroxyl of unmodified choline. Again, choline sulfate is known to be an osmolyte and protectant produced by certain fungi (*22*). Single-quadrupole mass spectrometry analysis of the fraction in which this compound elutes (fraction 115) revealed an m/z of 184.1, corresponding to protonated choline sulfate, further supporting its identification.

The third strongest peak in our mFL segment did have a match in the GISSMO database (arginine), but why should arginine, among all the common amino acids, be so abundant? Some searching among other identifiable spectral vectors produced a slightly modified version of ornithine (fraction 115) and a probable dimer of the modified ornithine and normal ornithine (fraction 128). Why make ornithine multimers? This, too, has an answer. Multimers of ornithine, modified to carry an N-oxide in place of an amide nitrogen, are the basis of a class of compounds called siderophores. These are iron-chelating compounds of interest in the fields of medicine, agriculture, and environmental decontamination (*23*). Searching for articles published in the last five years that cite “siderophore” yields more than 5,000 articles. So, metabolomics analysis of isolates from a variety of organisms can be interesting. The unexpected presence of certain compounds is a stark reminder that metabolite production can vary widely from one organism to another, reflecting the environmental conditions in which life forms must survive.

## MATERIALS AND METHODS

### Preprocessing and Peak Tracing

The metabolite fraction library (mFL) was imported into MATLAB, and preprocessing was performed, including the removal of solvent regions such as water and formic acid, which was a component in the mobile phase of the HPLC used to make the library (*11*). Initial peak picking was done fraction by fraction using a user-defined intensity threshold. Identified peaks were traced longitudinally across fractions to map elution profiles. A peak was sequentially linked to an active profile if it appeared within a strict 0.003 ppm margin of the preceding fraction’s peak location. The profile was terminated when no peak was detected within this chemical shift tolerance. To improve correlation, zero-intensity values were padded to the beginning and end of each bounded profile. Finally, to eliminate redundant profiles, those that span the same set of fractions were compared; if their Pearson correlation exceeded 0.99, only the profile with the highest absolute intensity was retained.

### Target-Driven SVD and Alternating Least Squares (ALS) Optimization

For each isolated profile, SVD was executed on a sub-matrix containing only the fractions where that specific profile elutes. This localized SVD decomposed data into three distinct matrices: U (elution profiles), S (singular values), and V (spectral vectors). An Alternating Least Squares (ALS) optimization loop was used to find optimal rotation matrices that best linearly recombine the abstract SVD components into physically meaningful elution and spectral vectors. Crucially, each optimization was designed to extract and isolate only a single target feature. The first principal component of the SVD matrices acted as the mathematical starting point for the optimizer; specifically, the first vector of U initialized the elution optimization, while the first vector of V initialized the spectral optimization. While the algorithm solves exclusively for this primary component, the remaining SVD components were incorporated into the calculation as a mathematical sink, absorbing noise and overlapping signals.

The elution profiles were optimized first. The solver minimized the difference between the traced elution profile and the first component of U, subject to non-negativity and L2 normalization. To guide spectral optimization, a target covariance spectrum was generated by computing correlations between the traced elution profile and all chemical shifts in the localized fraction sub-matrix. Peaks exhibiting a correlation below 0.97 to the elution shape were zeroed, as were any signals falling below 10% of the maximum covariance intensity. The resulting target was renormalized to 1. The spectral optimization minimized the deviation from the target covariance spectrum while enforcing non-negativity, targeted sparsity (which heavily penalizes signal in regions with low correlation), and a Tikhonov tethering penalty that forces the elution and spectral weight matrices to converge. The algorithm iteratively alternated between optimizing the spectral weights and the elution weights. The loop terminated when the weight matrix changed by less than 10% between iterations, or after a maximum of 10 iterations.

### Spectral Cleanup

Upon ALS convergence, the elution and spectral profiles were reconstructed from the original SVD matrices using the optimized weights. The elution profile was extracted by matrix multiplication of the full U matrix by the elution rotation matrix. The spectral profile was extracted by left-dividing the spectral rotation matrix by the full V matrix. Finally, the transformed S matrix was obtained by left-dividing the elution rotation matrix by the product of the full S matrix and the spectral rotation matrix. The extracted elution and spectral profiles were then further cleaned. For the elution profile, any negative elution values were set to zero. For the spectral profile, peaks that were highly correlated with the elution profile (greater than 0.90) were subjected to a baseline intensity cutoff of 5% relative to the maximum peak height. Moderately correlated peaks (0.70 to 0.90) faced an ascending penalty gradient, capping at a maximum 15% intensity cutoff for peaks at the 0.70 threshold. Both the elution and spectral profiles were normalized after the cleaning step. Before continuing the workflow, a quality-control measure was performed where any purified spectrum that showed a Pearson correlation below 0.2 with its initial target covariance spectrum was permanently discarded.

### Hierarchical Clustering and Global Error Minimization

To consolidate redundant components representing the same underlying metabolite, the purified spectra were compared using a pairwise Pearson correlation matrix and grouped by complete-linkage hierarchical clustering. The algorithm performed a sweep over correlation cutoffs (ranging from 0.50 to 0.95) to iteratively prune the hierarchical tree and assess the resulting configurations.

For each cutoff tested, a consensus spectrum was first generated for each cluster by computing the point-by-point median of its constituent spectra. Next, to determine the optimal elution profile for each cluster, the algorithm employed an iterative selection loop. Rather than rely on averaged elution profiles, the algorithm systematically tested each member’s exact physical elution profile. It substituted a member’s physical profile into a global elution matrix, multiplied this matrix by the consensus spectra to simulate a fully reconstructed fraction library, and calculated the Frobenius norm of the difference against the original experimental library. For each cluster, the algorithm retained the specific physical elution profile that minimized this global reconstruction error.

Once the optimal elutions and consensus spectra were paired, the clusters were ranked by a composite purity score that evaluated target-covariance match, intra-cluster similarity, and peak intensity. The algorithm then incrementally added these ranked components to the model one by one to find the optimal subset size (the number of metabolite spectral vectors, **k**) that minimized the relative reconstruction error. Ultimately, the final correlation cutoff was selected based on which parameter combination and component count yielded the lowest Frobenius-norm error relative to the original fraction library.

### Integration

Automated integration via SVD was achieved by computing the outer product of the purified elution and spectral vectors to determine the total integrated volume for each isolated metabolite across the fraction library. This volume was compared to the integrated volume of the internal standard, DSS, across the corresponding fractions. Finally, metabolite concentrations were calculated using the ratio of total integrable protons in the isolated spectrum relative to the DSS reference protons.

### Database Search

The final step in fraction library analysis was compound identification. This was accomplished by comparing our extracted spectra to spectra of known metabolites. At the time of our application, spectra of common metabolites were available in several databases, including the Biological Magnetic Resonance Database (BMRB), the Human Metabolite Database (HMDB), and the GISSMO database (entries derived from BMRB). The advantage of the latter database is that spectra acquired (or simulated) at all common magnetic field strengths are available. This is also true for many entries in the HMDB. ^1^H NMR spectra are composed of multiplets, generated by scalar coupling of spins; for first-order spectra, these are centered at the chemical shift of each chemically distinct proton. The chemical shifts, on a frequency scale, are linearly dependent on magnetic field strength, while scalar couplings are not. Moreover, multiplet structure varies as the chemical-shift difference between coupled protons approaches the magnitude of the scalar coupling (strong coupling effects). These effects lead to spectra with complex field-dependent effects. Many NMR-based metabolic studies avoid this complexity by focusing only on chemical shift. However, this disregards important structural information found only in scalar couplings.

Hence, we preferred to directly compare an extracted spectrum with a database spectrum at the proper field strength (800 MHz in our case). We began with 1324 spectra at 800 MHz from the GISSMO database and added 6 spectra (mostly carbohydrates), beginning with data from the BMRB and literature searches. In these cases, we either found literature with complete lists of chemical shifts and scalar couplings (*16, 24*) or extracted the couplings and shifts by fitting spectra to ones generated by the simulation software, SPINACH (*17*). We then used SPINACH to produce accurate 800 MHz spectra from chemical-shift and coupling-constant lists. In each case, spectra were scaled to match the spectral size and chemical shift referencing of our fraction library. Database entries and our extracted spectra were also normalized to a value of 1.

Scoring of extracted spectra against database spectra was performed using a dot product between spectral pairs (since the spectra are normalized, this equals cosine similarity). For noiseless spectra, the score would be 1 for a perfect match and zero for no match. There is one complication that we can’t avoid. Some chemical shifts will vary with pH, and possibly other sample conditions. While efforts were made to select standard conditions (pH 7.0, 298 K, water as solvent in our case), it is not possible to include all effects in a database. Hence, we segmented the extracted spectra at points where multiplets were well-separated and iteratively offset segments by up to 0.02 ppm, retaining only the highest-scoring segment. For efficiency, in the present case, we worked with only two segments separated by the water resonance.

## Funding

National Science Foundation EEC-1648035

National Institutes of Health R35 GM145348

A pilot award from Georgia Tech and the Georgia Research Alliance

## Author contributions

Conceptualization: CE, JHP

Data curation: CE, JHP

Formal analysis: CE, JHP

Funding acquisition: ASE

Investigation: CE, JHP

Methodology: CE, JHP

Software: CE, JHP

Supervision: ASE, JHP

Validation: CE, JHP

Visualization: CE, JHP

Writing – original draft: CE, JHP

Writing – review & editing: CE, ASE, JHP

## Data, code, and materials availability

The *N. crassa* raw data: usnan.org/ark:/83454/c185a05632-633d-40c3-b39f-3b198a9ddd21.

The extended GISSMO Library at 800 MHz and SVD analysis code is available at https://github.com/edisonomics/Metabolite_Fraction_Library_SVD

## Supplementary Materials

This PDF file includes:

Figs. S1 to S2

Other Supplementary Material for this manuscript includes the following: Table S1

