## Supplementary material for "Singular Value Decomposition for the Automated Analysis of NMR Fraction Libraries": Figs. S1 & S2

**Supplementary Materials:**

This PDF file includes:

Figs. S1 to S2

Other Supplementary Material for this manuscript includes the following:

Table S1

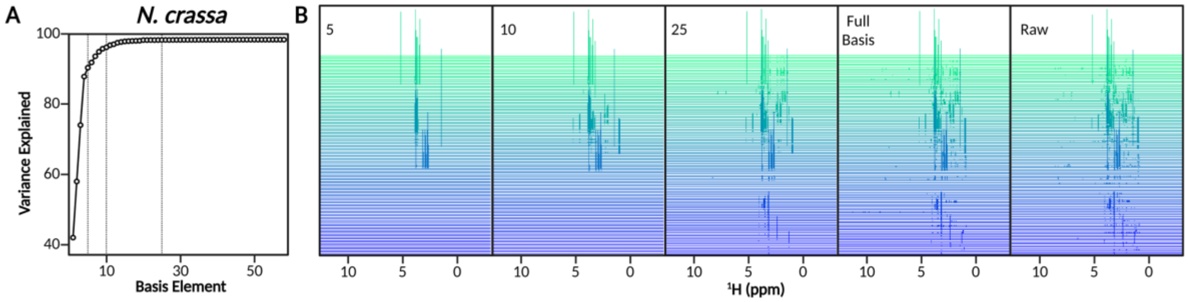


**Figure S1. Reconstructing the mFL** (A) Scree plot showing the increase in variance explained by adding additional purified basis set elements to the outer product calculation. Dashed vertical lines represent the number of basis set elements at 5, 10, and 25. The basis set elements are re-ordered by maximum elution intensity. (B) Visualization of the mFL as more basis set elements are added to the outer product calculation. The first three libraries show products from the top 5, 10, and 25 metabolite basis set elements. The full basis is the reconstruction using the full mBS of the first iteration of the SVD method. The raw library is the experimental library.

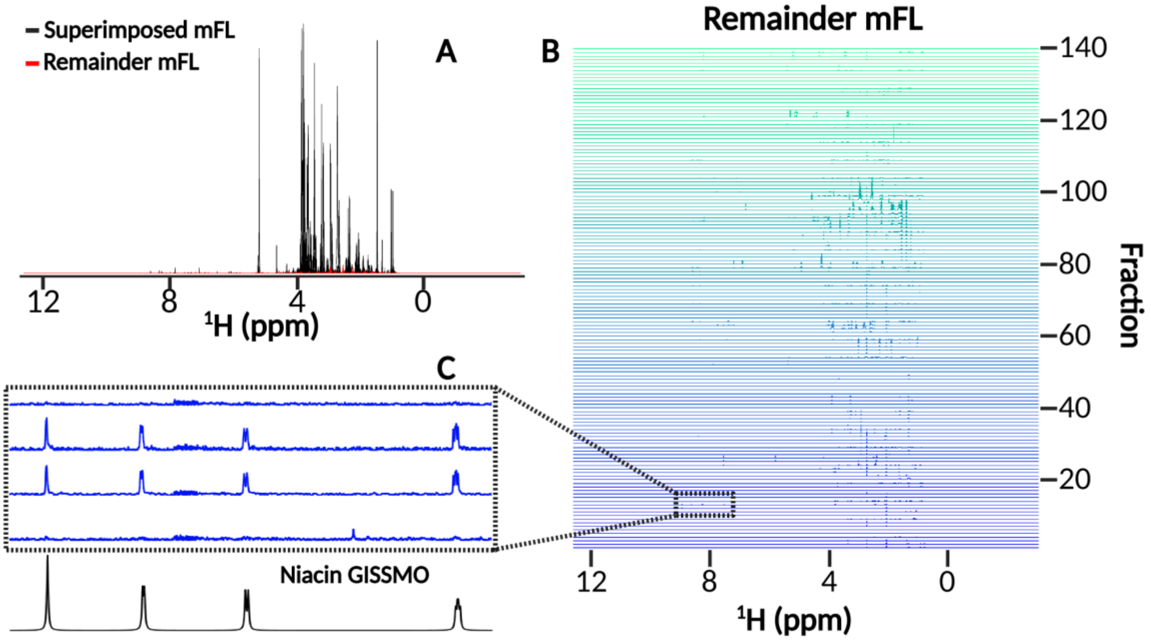


**Figure S2. Residual metabolites in the mFL.** (A) Every fraction of the *N. crassa* mFL (black) overlayed with the remainder of the same fractions after zeroing regions from mBS (red) (B) Visualization of the mFL after zeroing regions from mBS. (C) A zoom in of a low intensity region of the mFL showing peaks that match the Niacin database spectrum from GISSMO.
